# Accelerating Sequence Alignment to Graphs

**DOI:** 10.1101/651638

**Authors:** Chirag Jain, Alexander Dilthey, Sanchit Misra, Haowen Zhang, Srinivas Aluru

## Abstract

Aligning DNA sequences to an annotated reference is a key step for genotyping in biology. Recent scientific studies have demonstrated improved inference by aligning reads to a variation graph, i.e., a reference sequence augmented with known genetic variations. Given a variation graph in the form of a directed acyclic string graph, the sequence to graph alignment problem seeks to find the best matching path in the graph for an input query sequence. Solving this problem exactly using a sequential dynamic programming algorithm takes quadratic time in terms of the graph size and query length, making it difficult to scale to high throughput DNA sequencing data. In this work, we propose the first parallel algorithm for computing sequence to graph alignments that leverages multiple cores and single-instruction multiple-data (SIMD) operations. We take advantage of the available inter-task parallelism, and provide a novel blocked approach to compute the score matrix while ensuring high memory locality. Using a 48-core Intel Xeon Skylake processor, the proposed algorithm achieves peak performance of 317 billion cell updates per second (GCUPS), and demonstrates near linear weak and strong scaling on up to 48 cores. It delivers significant performance gains compared to existing algorithms, and results in run-time reduction from multiple days to three hours for the problem of optimally aligning high coverage long (PacBio/ONT) or short (Illumina) DNA reads to an MHC human variation graph containing 10 million vertices.

**Availability:** The implementation of our algorithm is available at https://github.com/ParBLiSS/PaSGAL. Data sets used for evaluation are accessible using https://alurulab.cc.gatech.edu/PaSGAL.

## I. INTRODUCTION

Examining an individual’s genetic variations is typically carried out by mapping DNA fragments sequenced from the individual to an annotated reference. Accuracy of this process is critical to draw correct conclusions in biological and medical scenarios [1]. The traditional practice in genomics has been to represent the reference as a sequence. Linear representation of a genome can however, only capture a single individual (in fact, a single haplotype copy within the individual), and therefore does not use the extensive genomics data now available across multiple individuals and populations. Recent biological studies have demonstrated superior genotyping accuracy in variant-rich regions of the human genome by replacing the reference sequence with a variation graph, i.e., a reference sequence augmented with known genetic variations [2]–[5]. The graph-based representations of the reference sequences are increasingly gaining traction due to their natural ability to succinctly represent population-wide variations [6]–[8]. Continued improvements in sequence to sequence alignment algorithms were pivotal to establish it as a fundamental routine for measuring evolutionary distance in biology [9]. Similarly, fast and accurate sequence to graph aligners are required to fully realize the potential of graph-based reference representations.

Broadly, the variation graph is a directed graph with vertices or edges labeled with DNA sequences. Following prior works [10]–[13], we represent the variation graph as a directed acyclic graph (DAG), with each vertex labelled with a DNA base. Standard dynamic programming (DP) based sequence to sequence alignment algorithms [14] can be extended to work with DAGs [15]. For an input DAG *G*(*V, E*) and a query sequence of length *m*, sequence to DAG alignment problem can be solved sequentially in *O*(*m*(|*V*|+|*E*|)) time [15], [16].

The sequence to DAG alignment problem becomes highly compute-intensive on real input data sets. The variation graphs associated with some of the most diverse regions (e.g., MHC, LRC segments) in the human genome contain vertices and edges in the order of millions. Moreover, we are typically required to solve several instances of this problem for each query read in high-throughput sequencing data sets, further adding to the computational complexity. As a consequence, time to solution using a naive sequential algorithm would require multiple days or months. To resolve this computational bottleneck, the focus of this work is to develop the first parallel algorithm that takes full advantage of the modern wide SIMD multi-core architectures.

Different sequencing technologies produce reads of varying characteristics. Normally, the fixed length (100 or 150 bp) reads from Illumina sequencers are referred to as short-reads, whereas the much longer variable length reads produced by PacBio and ONT sequencers are referred to as long reads (mean length >10 Kbp). Several seed-and-extend based heuristic algorithms have been recently proposed for solving the alignment problem that can handle large data sets [8], [13], [17]–[20]. These algorithms so far have mainly focused on short read mapping. Typically, such algorithms employ an index based approach to quickly narrow down the search space during the alignment process. In particular, substrings that span all possible alternatives in the graph are indexed using classic string data structures. Due to the exponentially growing potential number of paths as a function of number of variants, existing heuristics do not translate into a practical solution for dense variant-rich graph regions or aligning long reads [21]. We seek to remedy this by accelerating the exact DP algorithm which is suitable for both long and short read data, and complex graphs.

In this work, we develop a three-stage parallel approach to accelerate the dynamic-programming based sequential algorithm, where the first two stages compute the two end points of an optimal alignment, and the last stage executes a traceback procedure to compute base-to-base alignments. Each of these stages leverages inter-task parallelism such that multiple reads are processed independently in parallel. In addition, we propose a new blocked strategy to compute the DP score matrix that ensures high memory locality, thus allowing us to efficiently utilize wide SIMD width and multiple cores without exceeding constraints imposed by memory bandwidth. These optimizations enable near-linear weak and strong scaling behavior of the algorithm using 48 cores. As a result, we are able to align long and short MHC read sets with 10x coverage to MHC variation graphs in about three hours or less, which would otherwise take multiple days to process. Finally, we show superior performance in terms of runtime and accuracy against existing exact and heuristic algorithms, respectively.

## II. BACKGROUND

### A. The Sequence to Graph Alignment Problem

Although the primary motivation for this work lies in the domain of biological data analysis, string algorithms for approximate matching to graphs have been actively studied for the past three decades for different application areas, e.g., text retrieval, matching regular expressions, and pattern matching on hypertext [16], [22]–[24].

The classic sequence to sequence alignment problems for approximate matching are typically classified as either global, semi-global or local alignment. These problems can be solved exactly using dynamic programming (e.g., using Needleman-Wunsh [25] or Smith-Waterman [14] algorithms). Given a scoring scheme to reward matches and penalize mismatches, insertions and deletions, the alignment problems are formulated to compute alignments that achieve maximum score. Similarly, when aligning a query sequence to a variation graph, the problem is to identify the highest scoring alignment between the query sequence and any path in the graph.

In this work, we focus on the sequence to DAG alignment problem in local mode, i.e., computing local regions of similarity [14]. The proposed parallelization algorithm in this paper generalizes to other alignment modes, but they are not discussed for brevity. Following previous works [2], [10]– [13], we consider variation graph as a DAG *G*(*V, E, σ*), where function *σ* assigns each vertex a character from the alphabet set Σ = {*A, C, G, T*} describing DNA bases. Naturally, any path *p* in the graph spells a DNA sequence. Let *q ∈* Σ^*∗*^ be a query sequence of length *m*.

#### Definition

Sequence to DAG Local Alignment Problem: Given a query sequence q and a DAG G(V, E, σ), identify a path p in the DAG and a substring of *q*: *q*[*i..j*] s.t. the optimal alignment score between *q*[*i..j*] and the sequence specified by *p* is maximum over all possible choices for *p*, *i*, and *j*. In addition, report the corresponding alignment.

In cases when the query has multiple optimal alignments, we aim to output one of them. Although the problem definition includes a single query sequence for convenience, we are required to solve numerous instances of the problem, twice for each input sequence (counting both the complementary DNA strands) in the set of reads being mapped.

### B. Sequential Algorithm

Sequence alignment to DAGs is computed using dynamic programming (DP), essentially by extending the Smith-Waterman algorithm to DAGs [15], [16]. Assume the DAG *G* is topologically sorted. Suppose *C*_*i,j*_ denotes the highest score of an optimal alignment between any suffix of *q*[1..*i*] and any path ending at vertex *v*_*j*_. Then, a sequential *O*(*m*(|*V*|+|*E*|)) time algorithm follows from the recurrence below:

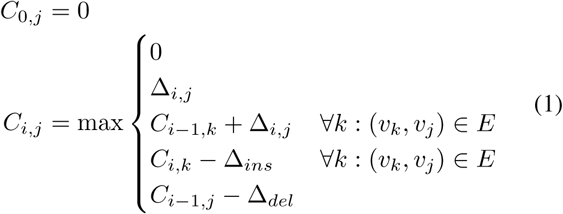

where Δ_*i,j*_ denotes the score of a match/mismatch, and Δ_*ins*_ and Δ_*del*_ denote the insertion and deletion penalties, respectively. The DP score matrix has a height of *m* + 1 and width of |*V*|. Note that score of each cell depends on cells in the previous row as well as cells to the left in the same row. Once the location of optimal alignment in the matrix is known, final base-to-base alignment is reported using a traceback procedure that follows the chain of decisions made in computing the *C*_*i,j*_’s. Similar to the Smith-Waterman algorithm, the traceback begins at the highest scoring cell and proceeds until a cell with zero score is encountered. The two cells with the zero and the highest score denote the start and the end of an optimal alignment, respectively.

### C. Constraints on Design of Parallel Algorithm

The described sequential algorithm is similar to the Smith-Waterman algorithm, the only difference being that each vertex can now have multiple neighbor vertices instead of just one (Figure 2). This one difference, however, makes numerous parallelization strategies [26]–[29] developed for accelerating the Smith-Waterman algorithm either inapplicable or inefficient for the sequence to DAG alignment problem. We list the challenges below:

**Fig. 1:**
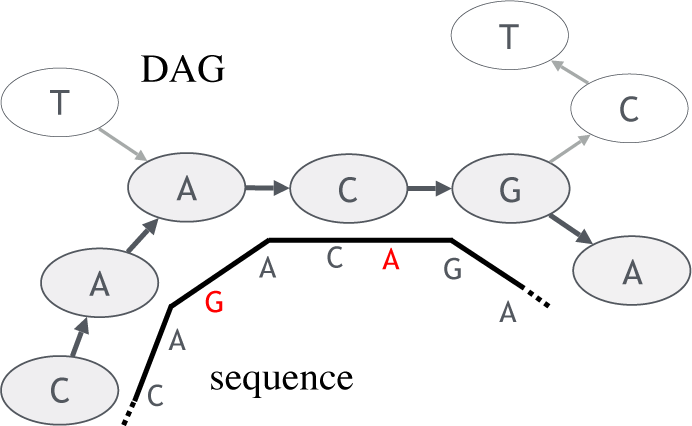
An example of an alignment of a sequence along a path in a DAG, while accounting for sequence errors (denoted in red).

**Fig. 2:**
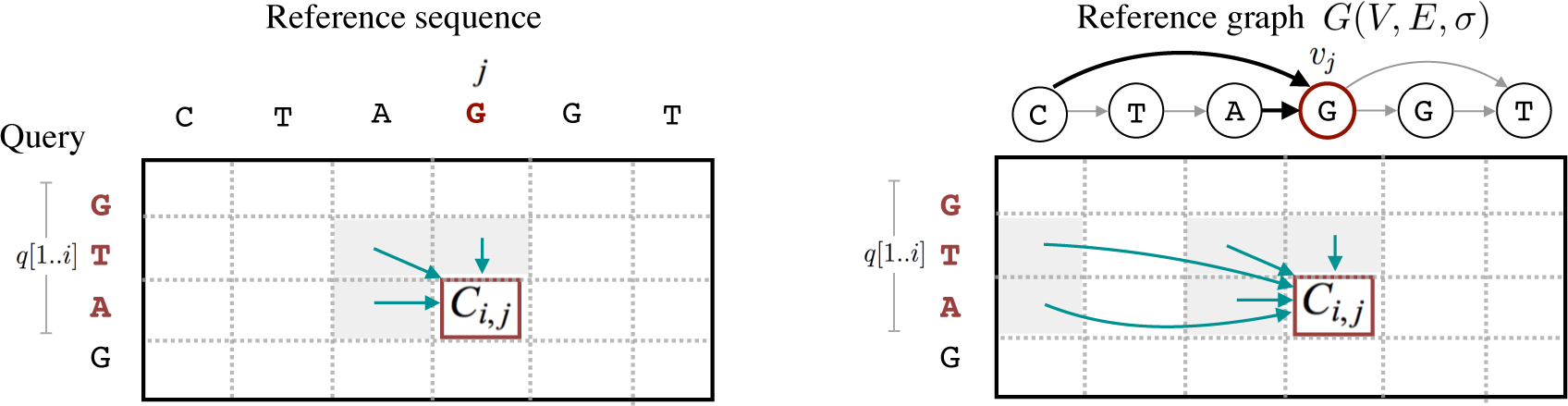
Example to illustrate difference between Smith-Waterman sequence to sequence alignment and sequence to DAG alignment procedures.

1. Storing complete DP score matrix in memory is usually impossible with real input data. In the Smith-Waterman algorithm, score matrix can be computed either one column, row, or diagonal at a time. This is possible because storing one previous column (or row) is sufficient as the DP progresses. However, vertices in the variation graph can be connected to many (near and distant in topological order) predecessor vertices, leaving row-wise computation as the only choice.
2. Unlike the Smith-Waterman algorithm, count of arithmetic operations required to compute score for each cell *C*_*i,j*_ in a row is not uniform, and depends on in-degree of vertex *v*_*j*_. This further makes SIMD-based intra-task parallelization challenging.

We present a new inter-task based parallelization approach that takes into account the above constraints.

## III. PROPOSED PARALLEL ALGORITHM

### A. Graph Representation

Solving Recurrence (1) requires frequent access to graph vertices and edges. Therefore, it is important that we spend as few CPU cycles as possible to access graph information while computing scores. Variation graphs are highly sparse, in fact the edge to vertex ratio is typically close to one [7]. Therefore, we choose the standard ‘compressed sparse row’ (CSR) format. It allows constant time access to adjacency list of any vertex. In this format, we use three arrays: the first one of size |*E*| for contiguous storage of adjacency list, another (|*V*|+1)-sized pointer array to mark start and end offsets for each vertex within the adjacency list, and the last array of size |*V*| to store DNA character labels of each vertex.

### B. A Three-Stage Algorithm

The proposed algorithm is designed to produce not only the optimal alignment scores, but also the base-to-base alignments corresponding to them. The base-to-base alignments are computed using a traceback procedure which requires access to the entire score matrix, or an appropriate section of it. In most practical cases, the sizes of the score matrices are too large to be able to completely store in memory. Therefore, we execute a three-stage algorithm to keep the memory-usage low. The first two stages of the algorithm are executed to identify starting and ending positions of the optimal alignments. In particular, the first stage *DP-fwd* computes ending position of an optimal alignment for each read by executing the DP to solve Recurrence (1). The second stage *DP-rev* solves the same DP in reverse direction, i.e., from bottom to top to locate the starting positions of the optimal alignments. As score of a cell depends only on its current and previous row (Section II), we only need to keep two rows in memory. Hence, the two stages use less memory. Finally, the third stage uses the starting and ending locations to recompute the corresponding section of the score matrix, and executes a traceback to report the base-to-base alignments (Section III-D). This approach is similar to the one proposed by Huang *et al.* [30] to compute local alignment between two sequences.

The first DP-fwd stage returns optimal score, ending position of an optimal alignment, and optimally aligning strand of each input read. DP-rev stage only uses sequences corresponding to the optimally aligning read strands as its input. Figure 3 summarizes the role of all three stages in the algorithm. Note that both DP-fwd and DP-rev stages compute the entire DP matrix using the same recurrence relation, therefore designing a single parallel strategy suffices to accelerate them. We propose a parallel algorithm in the following section.

**Fig. 3:**
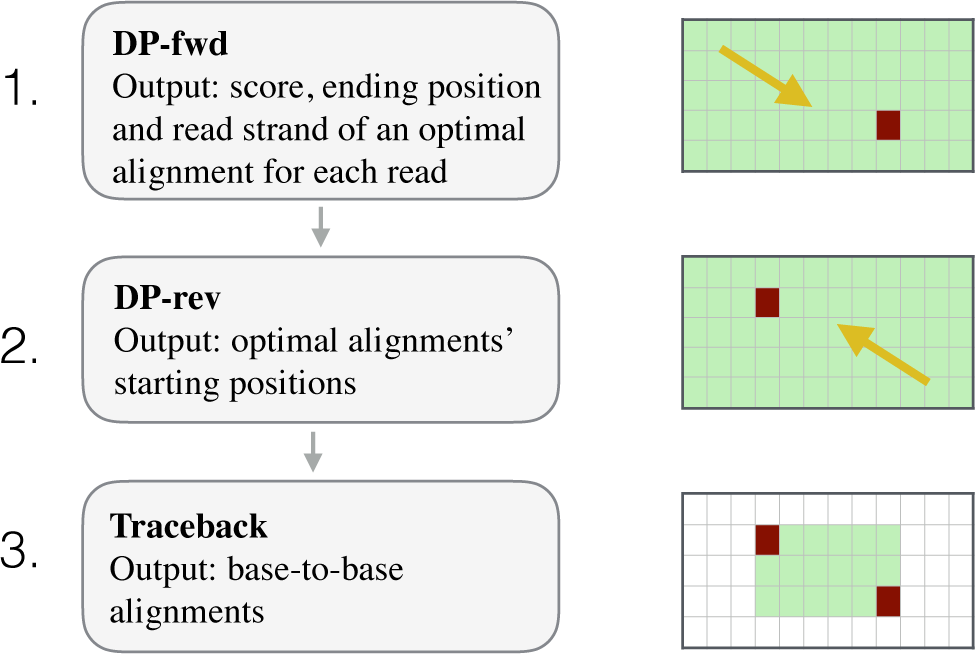
Role of the three stages used in our algorithm.

### C. Parallel Computation of the Score Matrix

Prior to describing full details, we give an overview of the algorithm. Both DP-fwd and DP-rev stages compute the entire score matrix containing (*m* + 1) × |*V*| elements for each input sequence. Computing score matrices is highly compute-intensive, and consumes most of the time in sequential as well as our parallel algorithm. The proposed algorithm is inspired from previous optimization efforts targeted towards accelerating Smith-Waterman alignment using SIMD instructions [31], [32]. Alignment of a single sequence is called a task. We present an inter-task parallel algorithm to accelerate the matrix computation. In other words, rather than parallelizing the alignment of a single read, the algorithm processes multiple reads simultaneously (Section III-C1). To compute each task, we can choose to follow a naive sequential algorithm, essentially computing the scores row by row. However, it turns out that traversing *O*(|*V*|)-sized row buffers repeatedly makes the algorithm memory-bound (Section III-C2). To address this issue, we subsequently introduce a new blocking strategy that leverages a domain-specific property of variation graphs, and enhances memory access locality.

#### 1) Inter-task parallelism

Our algorithm leverages inter-task parallelism by aligning multiple reads simultaneously. Below, we discuss how to make use of multiple threads and SIMD instructions to realize this efficiently:

##### Multi-threading

We divide the input read set into batches that are individually scheduled to different threads. As the runtime to align reads of different lengths varies, we leverage dynamic scheduling policy in OpenMP.

##### Vectorization

Within each thread, we vectorize our implementation to process all reads in a batch simultaneously (Figure 4a). Count of reads in a single batch is set to SIMD width to keep all vector lanes busy. For instance, recent Intel^®^ Xeon^®^ Skylake processors^1^ support AVX512 integer instructions (512 bit vectors). Therefore, depending on the requirement of precision to compute scores (e.g., int8, int16 or int32), there is scope of 16-64x speedup using vectorization. For each batch of reads, we convert read characters from AoS to SoA format to ensure that we can load the read characters for one cell update using just one vector load instruction. Suppose in-degree of vertex *v*_*j*_ is *δ*_*j*_, then computing *C*_*i,j*_ across all vector lanes uses 10 + 4*δ*_*j*_ vector operations (using cmpeq, blend, set, max, and add; not counting load and store) in our implementation. Finally, because read lengths within a batch of reads can vary, we pad shorter reads with dummy characters to obtain uniform lengths.

**Fig. 4:**
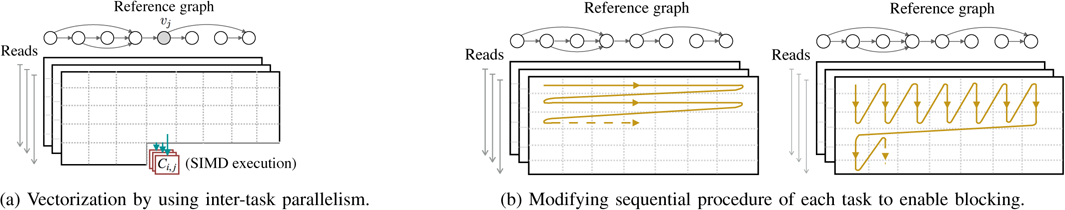
Visualization of the proposed parallel algorithm.

##### Load balancing

Lengths of long reads tend to vary significantly in a single sequencing run. Therefore, to avoid wasteful work due to padding, we sort the complete input read set by their length before dividing them into batches. In this way, variation in the lengths of adjacent set of reads is reduced. The sorting is done in decreasing order of read lengths to make sure that processing of longer reads is initiated first.

##### Optimizing Precision

Operating at lower precision (e.g., int16 vs. int32) yields higher scope for parallelism using vector units. Note that the product of maximum input read length and match parameter is an upper bound on the score value in all DP matrices. Based on this value, the algorithm decides the required precision at runtime. Besides maximum score, we also keep track of its column and row position during DP computation. The row positions use the same precision as score values because they are bounded by maximum read length. The column positions, however, can range from 1 to |*V*|, therefore they are always operated using int32 precision.

In the above inter-task parallelization scheme, each individual task can still be executed using a naive sequential algorithm, essentially computing the scores row by row. Working independently on individual reads gives us an advantage that there is no synchronization needed across threads or vector-units, which favors both performance and programmability. For each task, we need to maintain two score buffers for the current and previous rows. The two buffers can be used inter-changeably, i.e., one for reading previous row and one for computing current row. However, each of them uses *O*(|*V*|) memory, and does not fit in cache. As shown later in results (Section IV), this issue limits scalability by making the algorithm memory bandwidth bound due to frequent access to DRAM. We next describe a blocked computation strategy which modifies the sequential procedure of each task to resolve this issue.

#### 2) Improving Memory Locality using Blocked Computation

We propose a blocked algorithm to compute the score matrix which significantly reduces the average count of reads and writes to DRAM. The first step is to increase the granularity by computing multiple rows rather than one row in a single horizontal sweep (Figure 4b). For a subsequent horizontal sweep, we just need to preserve scores associated with the last row of the current block. Next, while processing multiple rows in a single horizontal sweep, we modify memory access pattern to ensure majority of accesses are cached. This modification leverages a domain-specific topological property of the graphs.

In the variation graphs, we find that the fraction of vertices connected to ‘distant’ vertices in the topological order is significantly small. More formally, let *B*_*width*_ be an appropriately chosen distance threshold, then the number of vertices in set *V′* = {*v*_*i*_: (*v*_*i*_, *v*_*j*_) ∈ *E, j* − *i* ≥ *B*_*width*_} is much smaller compared to |*V*|, for even small values of *B*_*width*_. In our implementation for instance, we selected *B*_*width*_ = 8 as this value was appropriate for various graphs tested empirically. This particular graph property is attributed to the fact that > 99.9% of genetic variations in a human genome are either single nucleotide substitutions or small insertions/deletions [33]. Such genetic variants mostly appear as small bubbles in the variation graphs. Large structural variants which would result in connecting farther vertices occur at much less frequency. As a result, majority of vertices in variation graphs are expected to have all their neighbors in near vicinity in the topological order. We next show how to leverage this property to improve the memory access pattern.

In the blocked-procedure, suppose the count of rows processed in a single horizontal sweep is denoted as *B*_*height*_. We use a small circular buffer of size *B*_*height*_ *· B*_*width*_ for temporary storage of scores while processing the *B*_*height*_ rows (see Figure 5). Using this buffer, score of a vertex *v*_*i*_ is available while computing score of *v*_*j*_ whenever *j - i* < *B*_*width*_. This modification ensures that majority of DRAM accesses are cached. To manage the scores of ‘long-hopped’ vertices ∈ *V ′*, we use a separate buffer to save their scores for subsequent access. This buffer is also small and manageable because |*V* ′| ≪ |*V*| . In our implementation, we set *B*_*height*_ and *B*_*width*_ to 16 and 8 respectively as these values resulted in the least memory latency and best performance during execution (further discussed later in Results section).

**Fig. 5:**
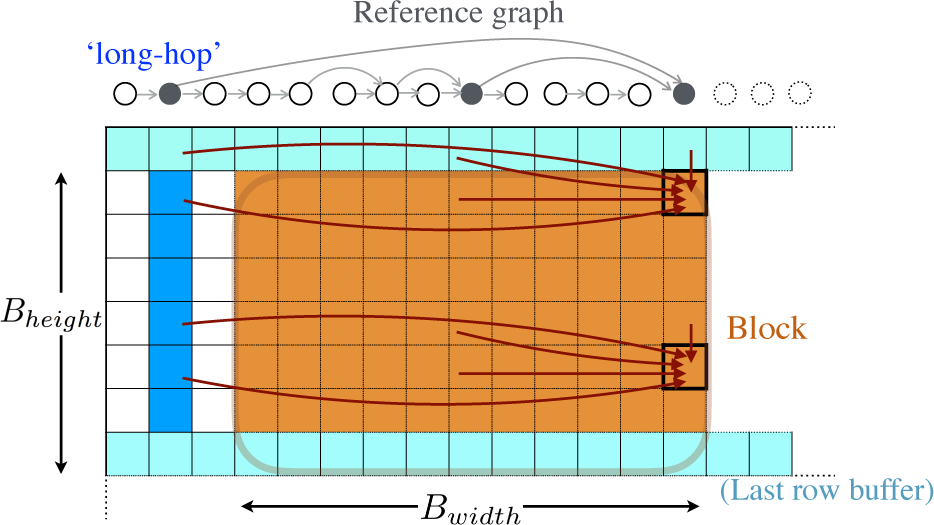
Visualizing a section of DP matrix to illustrate different memory accesses occurring using the blocked approach. Blocking improves memory locality because majority of accesses (red arrows) occur within the circular *B*_*width*_ *× B*_*height*_ block buffer.

### D. Computing Base-to-Base Alignments

The first two stages output the starting and ending alignment coordinates of each read. In the third stage, we recompute the enclosed section of the score matrix to execute the final traceback. Note that the coordinates of the optimal alignments of each read vary. Therefore, different matrix columns need to be computed for each read. This implies that we cannot re-use the inter-task SIMD parallelism that is developed for the first two stages. However, each of these tasks can still be executed independently using multi-threading. In this stage, we preserve the computed scores in memory. Because read alignments can hop through long edges (w.r.t. the topological vertex order) in the variation graph, even the score sub-matrices can be large. To reduce memory usage by a factor of four, we save the matrix using differences between adjacent rows instead. This helps because maximum absolute value of the differences (*C*_*i,j*_ − *C*_*i*-1,*j*_) is bounded by the sum of match and gap parameters [34], and can be saved as 8-bit integer values. We note that it is possible to extend alternate approaches to DAGs such as Hirschberg’s divide and conquer algorithm [35] or external memory algorithms [36] which use less memory, but they require more computation time in practice. Finally, the algorithm finishes after computing base-to-base alignments by tracing the paths associated with the respective optimal alignments of the input reads.

## IV. RESULTS

We refer to the C++ implementation of our **pa**rallel **s**equence to **g**raph **al**ignment algorithm as PaSGAL. This section provides details of the experimental setup, performance characteristics of PaSGAL, and advantages of the proposed optimizations. Subsequently, we demonstrate significant performance gains compared to existing read to graph aligners.

### A. Experimental Setup

#### 1) Data-sets

We used three input variation graphs and multiple read sets for evaluating PaSGAL (Table I).

**TABLE I:**
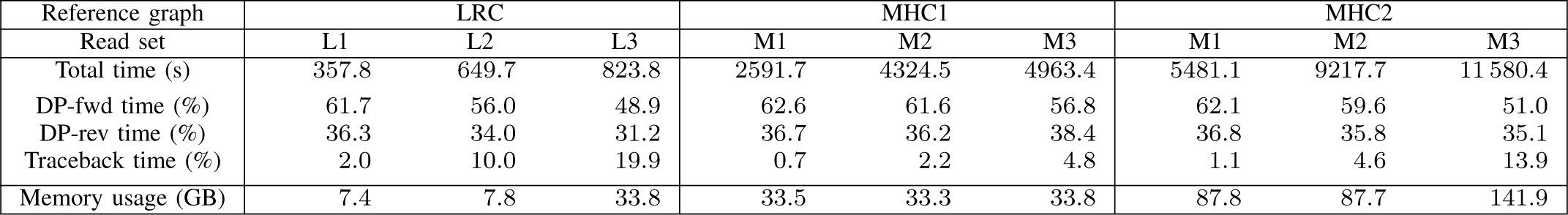
Summary of input variation graphs and read sets used for evaluation. First three columns show input graphs and their sizes while the remaining columns show characteristics of simulated read sets.

##### Variation Graphs

Leukocyte Receptor Complex (LRC) and Major Histocompatibility Complex (MHC) regions are among the most diverse variant hot-spots, spanning about 1.06 Mbp and 4.97 Mbp of the human genome [2], [7], respectively. We leveraged existing tools to build variation graphs using real public data. The first two graphs, labeled as LRC and MHC1 were built using the vg toolkit [8]. We supplied human genome (GRCh38) and variant files from 1000 Genomes Phase 3 [33] as input to vg. The variant files constitute small-scale variations (*≤* 50 bp) in genomes of 2504 individuals. To also evaluate using more complex graphs, we used a second MHC variation graph (MHC2) from a previous study [3], which also includes large structural variations.

##### Read Sets

Multiple sequencing read sets of different characteristics were simulated from the LRC (L1-L3) and MHC (M1-M3) regions in the human genome (GRCh38) (Table I). These read sets are representative of outputs produced using different technologies, lengths, and error characteristics. We used mason2 [37] and pbsim [38] tools to simulate single-end short Illumina and long noisy PacBio reads respectively. Sampling was done at 30x and 10x coverage from the LRC and MHC regions respectively. Because Illumina sequencers produce fixed-length short reads, L1 and M1 data sets contain uniform length reads of size 100 bp. On the contrary, long read technologies produce noisy reads of variable lengths; therefore L2-M2 and L3-M3 were sampled using mean read length 10 Kbp and 25 Kbp respectively. The minimum length, maximum length and mean error-rate parameters were set to 1 Kbp, 30 Kbp and 15% respectively during simulation. The match score, and mismatch, insertion, and deletion penalties were all set to 1. For each input read, the alignment program considers both complementary DNA strands, and outputs an optimal alignment to the input graph.

#### 2) Hardware/Software Description

Unless otherwise mentioned, all our experiments used a single node consisting of Intel^®^ Xeon^®^ Platinum 8160 (Skylake) processor in Stampede2 cluster located at Texas Advanced Computing Center. Each node is equipped with 192 GB RAM and two sockets, each containing 24 cores. Peak memory bandwidth of a node is 220 GB/s spread over two NUMA domains, one on each socket. These nodes operate at base frequency of 2.1 GHz, although frequency can vary due to turbo boost feature. Skylake platforms support AVX512 (512 bit) vector processing for 8-bit, 16-bit, and 32-bit integer operations.

PaSGAL was compiled using Intel^®^ compiler (v18.0.2). We used OpenMP for multi-threading and hand-written SIMD intrinsics for vectorization.

#### 3) Measurements

In all experiments, we measured runtime of the main alignment routine, and ignored pre-processing time, i.e., the time spent to load the input and converting graph into CSR format (Section III-A). Loading and pre-processing the input in PaSGAL took an insignificant fraction of time (*<* 1%). For all multi-threaded executions, we mapped a single thread to a single physical core.

### B. Performance Results

#### 1) Time to Solution

We first show the time to solution using PaSGAL for all nine input combinations in Table II using 48 threads. PaSGAL makes efficient use of multiple cores as well as vector units within each core to achieve fast time to solution. Using PaSGAL, we aligned 30x coverage read sets (L1-L3) to the LRC graph in *<* 15 minutes. For the larger MHC1 graph, 10x coverage read sets (M1-M3) were aligned in *<* 1.5 hours. Finally, the largest graph MHC2 took the longest time of 1.5 to 3.5 hours.

**TABLE II:**
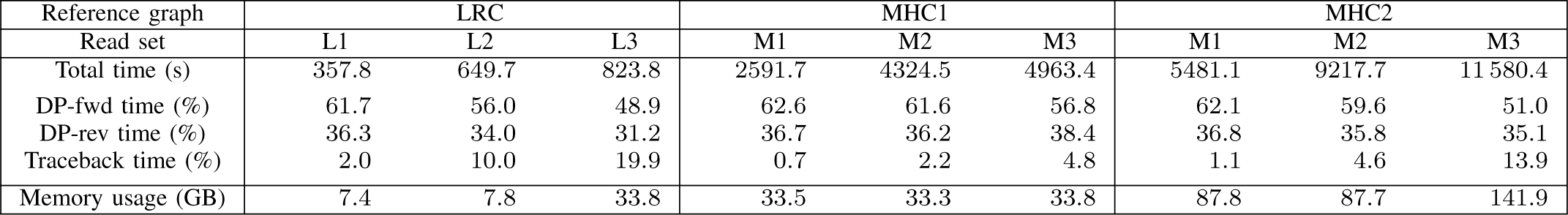
Performance evaluation of PaSGAL using all the test data sets.

We also show the performance achieved for score matrix computation as billion cell updates per second (GCUPS), the standard metric to evaluate Smith-Waterman algorithms (Figure 6). The GCUPS metric indicates the count of score matrix cells that are computed in a second, therefore higher is better. PaSGAL achieved peak performance of 317 GCUPS using the LRC/L1 input. To our knowledge, this is the highest performance achieved till date when aligning sequences to DAGs. Note that short read alignment (L1 and M1) was consistently fastest for all the three graphs because PaSGAL selects the required SIMD precision level based on the input read length (Section III-C1). For L1 and M1, 8-bit precision is sufficient as read length is only 100. On the other hand, the other read sets require 16-bit precision.

**Fig. 6:**
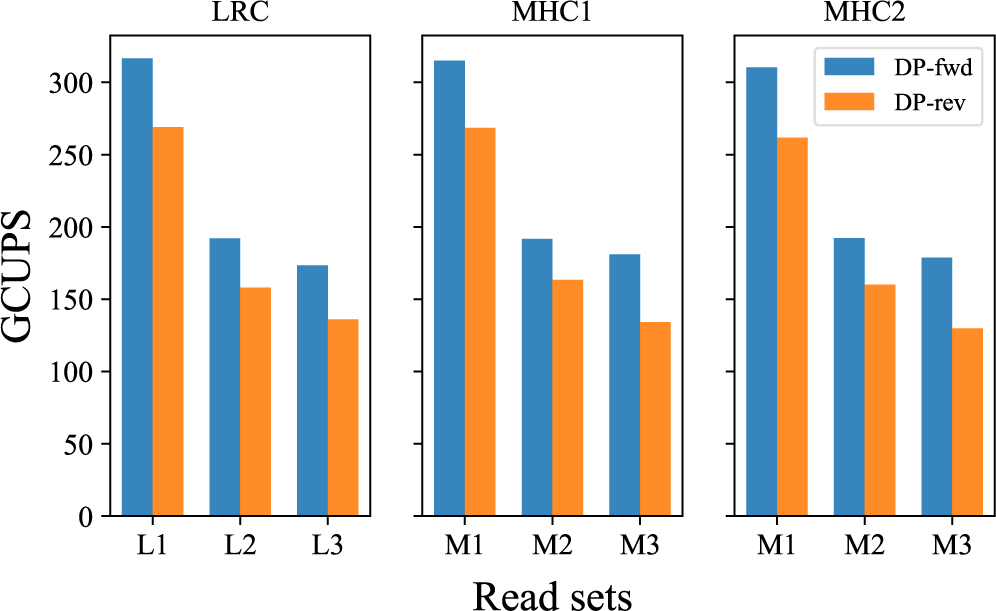
Performance achieved during DP-fwd and DP-rev stages of PaSGAL measured in billions of cell updates per second (GCUPS).

Large count of reads in an input set ensures that all the vector lanes do useful work, thus DP-fwd and DP-rev stages of the algorithm achieve high efficiency. We also note that the GCUPS performance numbers for forward DP are relatively higher compared to reverse DP. Even though the recurrences computed in DP-rev and DP-fwd stages are identical, DP-rev requires additional logic to ensure that it reports the end point of the same optimal alignment that was identified by the DP-fwd stage for each read. This is important because multiple alignments with maximum score can co-occur. Our algorithm guarantees to report one of those.

Finally, we also include a break-down of the total execution time into time spent in the three individual stages DP-fwd, DP-rev, and traceback (Table II). Even though the traceback phase is not vectorized, time to compute the two end points of optimal alignments using the DP-fwd and DP-rev stages still took majority of the time. This is because during the traceback stage, we are only required to compute a small portion of the score matrix.

#### 2) Load balance

Unlike short reads, long read lengths tend to vary significantly, therefore splitting the work equally can be challenging in an inter-task parallel approach. In PaSGAL, we address this issue by sorting the reads by their lengths and adopting dynamic scheduling policy (Section III-C1). We measured individual timings on 48 threads for all three stages of the algorithm. In Figure 7, we report ‘load imbalance ratio’ which is equal to maximum time divided by average time on all threads. Ideally, this ratio should equal one. We observe that this ratio is below 1.5 for all data sets. Better load balance is achieved for short read sets (L1, M1) relative to long reads, owing to their uniform lengths. Further, better load balance is observed for DP-fwd stage relative to DP-rev stage. DP-fwd stage processes both strands of DNA sequences, where as DP-rev only processes one, reducing its input size by half (Section III-B). The sorting approach is inherently more effective with higher read counts (Section III-C1).

**Fig. 7:**
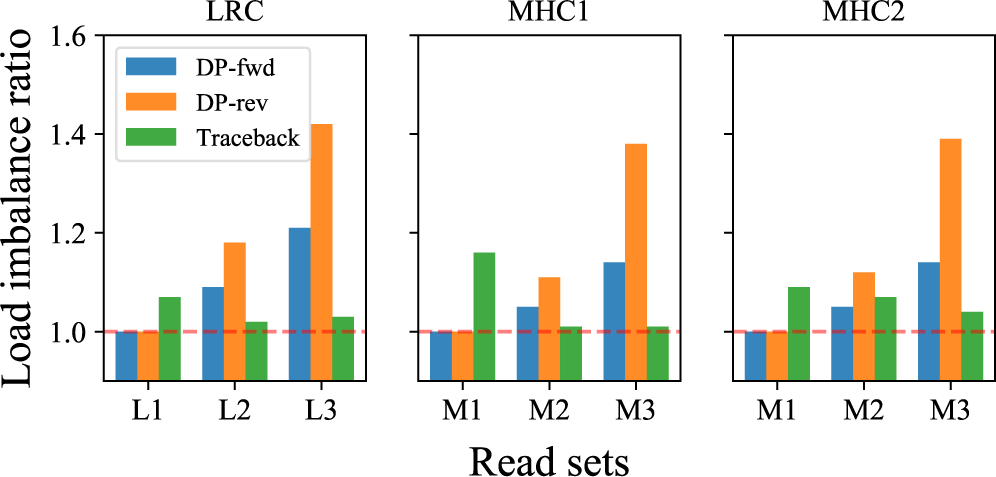
Load imbalance observed in PaSGAL using all test data sets while using 48 threads.

#### 3) Benefits of Proposed Optimizations

We next verified performance gains from the two optimizations – blocked strategy (Section III-C2) and vectorization (Section III-C1) used to accelerate the score matrix computation in DP-fwd and DP-rev stages, which account for majority of the time spent. We also collected critical performance counter numbers (e.g., memory latency, bandwidth etc.) using Intel^®^ VTune^®^ Amplifier tool. VTune profiling requires the experiments to be short to avoid counter overflow. Therefore, we used a small simulated short-read set from the LRC region, adding to 12,288 reads of length 96 bp, and aligned them to the LRC graph.

##### Blocked approach

Goal of the blocked approach is to remove memory access bottlenecks due to frequent access to DRAM (Section III-C2). PaSGAL uses default block width (*B*_*width*_) and height (*B*_*height*_) values of 8 and 16 respectively. Using 48 threads and vectorized execution, we analyzed benefit of this approach by manipulating block dimensions from 1 to 32 (see Table III). Note that using block height of one is equivalent to computing the score matrix one row at a time. The results support that the blocked approach succeeds in improving runtime by reducing memory latency, LLC (last-level cache) misses and DRAM bandwidth. This is because majority of reads and writes occur in the small *B*_*width*_ *× B*_*height*_-sized block. Each cell in the block is a SIMD register of size 64 bytes, therefore total storage memory equals 8 KB, making it small enough to fit in L1 cache memory (capacity=32 KB) of all cores.

**TABLE III:** Significance of using blocked approach.

| Block height | 1 | 4 | 8 | 16 | 32 |
| --- | --- | --- | --- | --- | --- |
| Time (s) | 40.9 | 11.1 | 8.4 | 7.8 | 8.0 |
| Avg. memory latency (cycle) | 27 | 8 | 7 | 7 | 8 |
| LLC miss count ($\times 10^6$) | 632.3 | 14.4 | 1.4 | 0 | 3.6 |
| Avg. DRAM bandwidth (GB/s) | 189.3 | 176.3 | 125.1 | 71.7 | 35.5 |

| Block width | 1 | 4 | 8 | 16 | 32 |
| --- | --- | --- | --- | --- | --- |
| Time (s) | 32.5 | 7.9 | 7.8 | 7.8 | 7.9 |
| Avg. memory latency (cycle) | 13 | 7 | 7 | 7 | 7 |
| LLC miss count ($\times 10^6$) | 44.6 | 0.5 | 0 | 0 | 0 |
| Avg. DRAM bandwidth (GB/s) | 177.2 | 69.8 | 69.6 | 70.8 | 70.6 |

(b) Performance analysis while increasing block width from 1 to 32, and keeping height fixed to the default value of 16.

##### Vectorization

Using Figure 8, we show the benefit of vectorization in PaSGAL by comparing it to our sequential scalar code, which computes score matrix sequentially in a row-wise manner. To isolate the benefit of vectorization and the enhanced memory locality, we executed this experiment using single thread only. Here we also experimented with three precision levels for integer instructions (8-bit, 16-bit and 32-bit). The plot shows that vectorization coupled with improved memory locality resulted in up to 58.7x speedup compared to the scalar code.

**Fig. 8:**
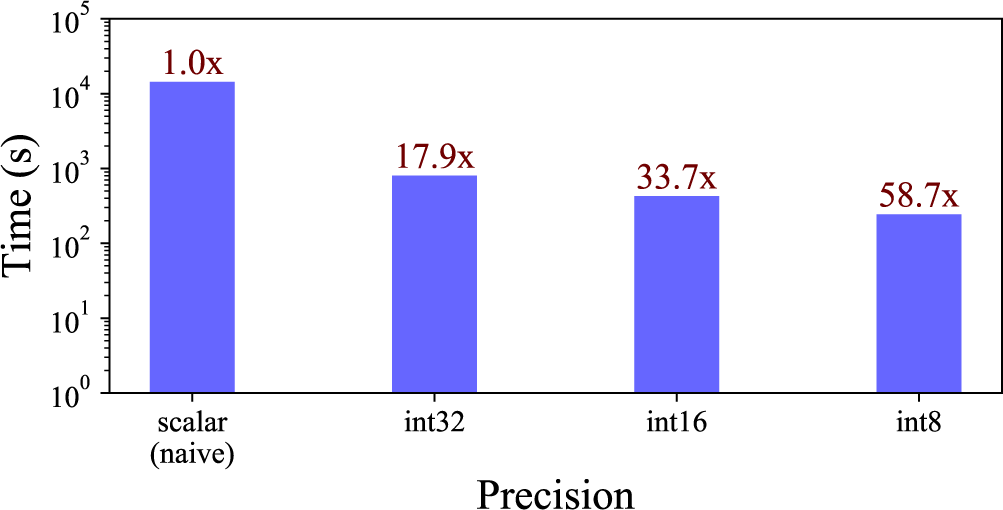
Performance improvement in PaSGAL obtained using improved memory locality and vectorization supporting different precision levels. Log scale is used for y-axis. This experiment was executed using a single thread.

#### 4) Scalability

The combination of our efficient vectorization strategy and blocking algorithm drive the high-performance in PaSGAL. These optimizations also helped us achieve near-linear strong scaling and weak scaling results going from 1 to 48 cores. The Skylake processors in Stampede2 use turbo technology, thus making scaling studies less reliable. Therefore, we conducted our scaling experiments on a different Skylake CPU (Intel^®^ Xeon^®^ Platinum 8180) with turbo technology disabled. In the two scaling experiments, we aligned L1-L3 read sets to the LRC graph, and report scaling behavior of the DP-fwd stage and overall runtime separately.

##### Strong scaling

Using the strong scaling experiment, we study the ability of our algorithm to compute the alignment problem faster with increasing core counts. Using 48 cores, the total runtime was reduced by 47x, 41x and 38x for the read sets L1, L2 and L3 respectively (Figure 9). Speedup factors for the DP-fwd stage were roughly similar-47x, 43x and 40x respectively. Short read set (L1), in particular delivered close to ideal scaling behavior because of the following two reasons-a) read count in L1 is very high, thus all SIMD lanes were busy doing useful work throughout the execution, and b) read lengths are uniform, therefore there was no overhead from load imbalance (Section IV-B2). We also evaluated the scaling behavior of the DP-fwd stage by manipulating the block dimensions (Figure 10). Results reveal that the blocked approach is critical for the near-linear speedups achieved.

**Fig. 9:**
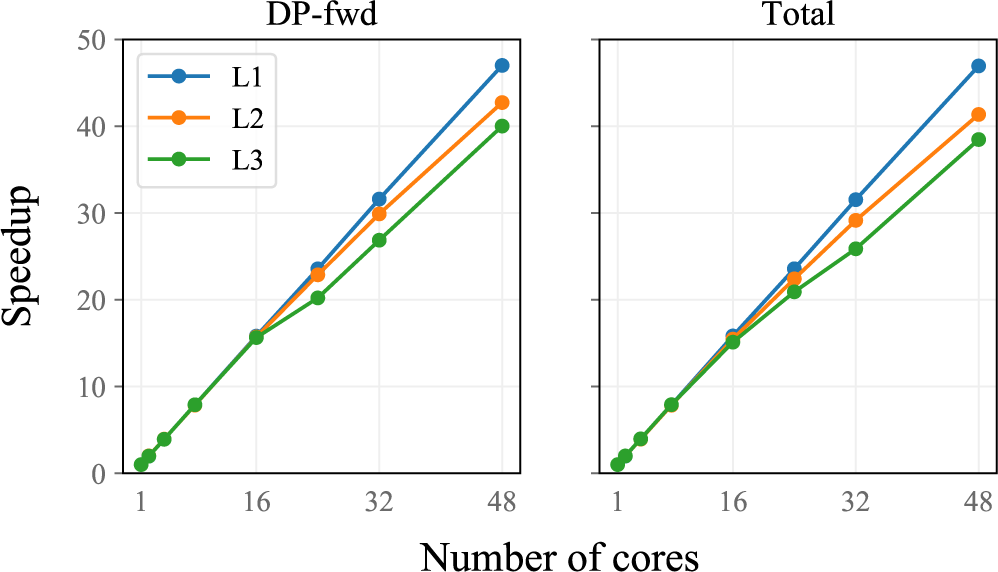
Strong scaling: Speedup achieved using PaSGAL with increasing core count relative to its single-core execution time. Left plot shows speedups achieved for the DP-fwd stage whereas right plot shows the overall speedup.

**Fig. 10:**
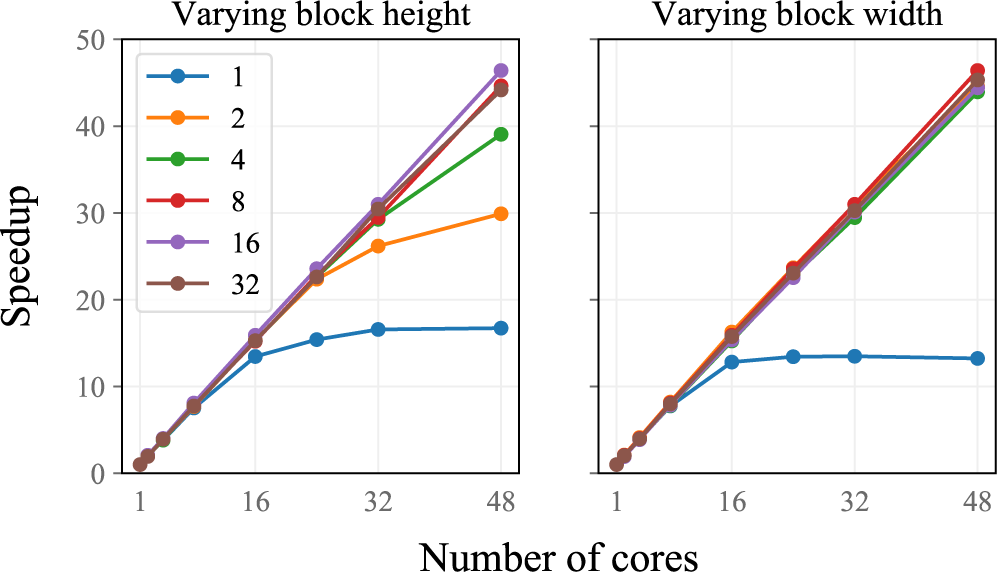
Illustration of how scaling behavior varies using different block sizes. Default values for block width and height were set to 8 and 16 respectively. We aligned the 96-bp short read set to the LRC graph in this experiment.

##### Weak scaling

In the weak scaling experiment, we maintained size of input read set proportional to core count. This metric measures the ability of an algorithm to handle larger input sizes given more resources. Therefore, an ideal weak scaling behavior translates to constant execution time regardless of core count. To conduct this experiment using 1 to 48 cores, we re-simulated read sets with coverage proportional to core counts (i.e., 30x for 48 cores, 15x for 24 cores etc.). Results show that nearly uniform runtime was achieved going from 1 to 48 cores (Figure 11).

**Fig. 11:**
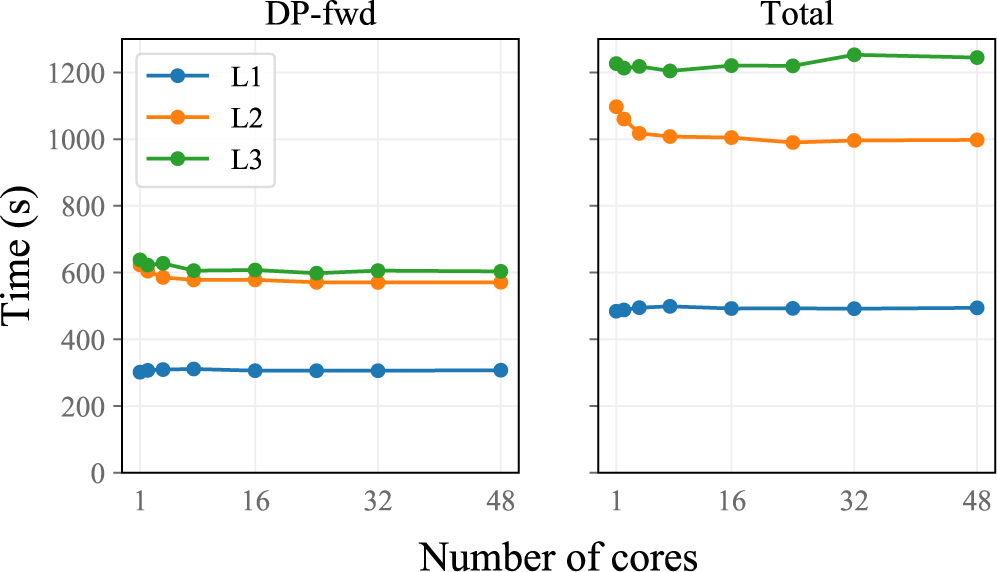
Weak scaling: PaSGAL’s execution time remains nearly uniform with larger input data and proportionally increased core counts.

### C. Comparison with Previous Algorithms

We compared PaSGAL against two recently published tools for sequence alignment to variation graphs – Graphaligner (commit:241565c) [39] and vg (v1.9.0-196) [8]. The software vg supports both exact and heuristic alignment modes. For convenience, we refer to its exact implementation as *vg-exact*, and heuristic implementation as *vg-heuristic*. We find that PaSGAL achieves up to 10x and 25x speedup against Graphaligner and vg-exact respectively, while using the lowest amount of memory. Compared to the vg-heuristic algorithm, we observe significant benefit in output quality. The comparisons against the exact and heuristic methods are discussed below separately.

#### 1) Comparison With Exact Algorithms

Graphaligner uses bit-level parallelism to compute edit distance between input reads and graph. The algorithm outputs edit distance scores, therefore we compared its runtime against the equivalent DP-fwd stage of our algorithm. We also utilized the sequential implementation available in the Graphaligner repository as our sequential baseline. vg-exact extends Farrar’s intra-task SIMD algorithm [40] to DAGs. It uses SSE (128-bit) intrinsics to accelerate the computation, and reports both optimal alignment scores and base-to-base alignment. As such, we compared its runtime against the total execution time of PaSGAL. Spoa [41], like vg-exact, also uses a intra-task SIMD parallelization algorithm for sequence alignment to DAGs, however, we could not compare against it because its implementation is designed to compute multiple sequence alignment for a different application. Both Graphaligner and vg-exact do not support multi-threading, therefore, we used a single thread for a fair comparison. To allow all the experiments to finish in reasonable time, we re-simulated six read sets: L1′-L3′, M1′-M3′ with 0.5x coverage by following the exact same procedure as before (Section IV-A1).

We show the speedups achieved using PaSGAL when compared to the other algorithms in Table IV. The speedups ranged from 40-98x, 3-11x and 13-25x when compared to the sequential baseline, Graphaligner and vg-exact, respectively. In three out of the six runs, vg-exact ran out of memory because it processes the DP column-wise and allocates the complete DP matrix in memory during the alignment. vg-exact supports affine gap penalty, which we plan to support in future versions of PaSGAL. Besides being fastest on a single-core, we conclude that PaSGAL also uses the lowest memory among all the algorithms.

Our previous results validate that PaSGAL supports efficient scalability using multiple cores (Section IV-B4), whereas current exact methods use single-thread only. Based on the observed numbers, it will take other algorithms multiple days to process the 10x coverage MHC read sets (M1-M3), which PaSGAL does in about three hours or less (Table II). Overall, PaSGAL provides significant advantage over existing exact algorithms in terms of its ability to process high throughput read sets and larger graphs.

#### 2) Comparison with Seed-and-Extend Based Heuristic Algorithm

PaSGAL being an exact aligner, guarantees to produce an optimal alignment, irrespective of graph topology and sequence characteristics. Heuristic algorithms accelerate the mapping process by avoiding a full-scale DP when mapping a read. The seed-and-extend based heuristics identify local graph regions using either maximal or fixed length exact matches, and validate the matches using an extension step. Although this approach can be significantly faster, both seed-computation and extension stages are still challenging to execute in dense variant-rich graph regions, and when the read lengths are long.

We ran vg-heuristic with default parameter settings using the L1′-L3′ read sets, and found it to be orders of magnitude (28-56x) faster than our exact algorithm. Next, we looked at the output accuracy. Since we executed both tools with the same scoring parameters, we compare the optimal alignment scores from PaSGAL against the scores computed by vg-heuristic (Table V). We note that a large fraction of output scores reported by vg-heuristic are sub-optimal for L2′ and M2′ read sets. It failed to produce reasonable output for L3′ indicating that its algorithm may be suitable for short reads only.

**TABLE IV:** Comparison with other exact algorithms.

| Reference graph | LRC |  |  | MHC1 |  |  |
| --- | --- | --- | --- | --- | --- | --- |
|  | L1’ | L2’ | L3’ | M1’ | M2’ | M3’ |
| vs. sequential | 94.9x | 50.3x | 41.1x | 98.3x | 56.3x | 51.7x |
| vs. Graphaligner | 10.7x | 4.2x | 3.0x | 10.0x | 3.7x | 3.7x |
| vs. *vg-exact* | 25.3x | 13.3x | - | 23.3x | - | - |

| Reference graph | LRC |  |  | MHC1 |  |  |
| --- | --- | --- | --- | --- | --- | --- |
|  | L1’ | L2’ | L3’ | M1’ | M2’ | M3’ |
| PaSGAL (GB) | 0.2 | 0.3 | 0.8 | 0.9 | 0.9 | 0.9 |
| sequential (GB) | 1.0 | 1.0 | 1.0 | 1.4 | 1.4 | 1.4 |
| Graphaligner (GB) | 1.1 | 1.1 | 1.1 | 2.0 | 2.0 | 2.0 |
| *vg-exact* (GB) | 0.7 | 108.8 | - | 2.9 | - | - |

(b) Peak memory-usage of all exact algorithms.

**TABLE V:** Accuracy evaluation of vg-heuristic algorithm for short and long read data sets. We compare its alignment score against the optimal score computed by PaSGAL.

| Read set | L1' | L2' | L3' |
| --- | --- | --- | --- |
| Fraction of alignments with<br>> 5% diff. from optimal score (%) | 0.04 | 24.53 | 100 |
| Fraction of alignments with<br>> 20% diff. from optimal score (%) | 0.00 | 9.40 | 100 |

## V. CONCLUSIONS

In this work, we presented an inter-task based parallel algorithm PaSGAL to accelerate alignment of sequences to DAGs. Although conceptually similar to the classic Smith-Waterman problem which admits easy parallelization, significant variability in the number and structure of dependencies in the dynamic programming table make parallelization of alignment to DAGs quite challenging. Given an input set of reads and a variation graph, PaSGAL outputs optimal alignment scores and base-to-base alignments, a requirement for downstream biological analysis. To the best of our knowledge, it is the first parallel algorithm for solving this problem that fully utilizes modern architectures by leveraging multiple cores and wide SIMD width. To achieve these goals, we presented a three-stage algorithm and several optimizations to maximize integer operations per second. As a result, we are able to compute alignments of high-coverage long or short read sets to large variation graphs associated with clinically important human genome segments in the order of few minutes or hours, which was not feasible with prior algorithms.

Besides pan-genomics, PaSGAL can be useful for other applications that benefit from sequence to DAG alignment, e.g., sequence alignment to splicing graphs in transcriptomics [42] and antibiotic resistance profiling [43]. Future work includes development of intra-task algorithms for use-cases with small count of query sequences (e.g., when aligning assembly contigs to graphs), and extending this framework to accelerate the alignment to general sequence graphs [44]. Our algorithm combined with an appropriate graph localization heuristic could scale to variation graphs of complete vertebrae genomes. There is plenty of evidence in recent scientific literature that justifies the utility of variation graphs as a reference for studying genetic variants. The scalable and exact approach presented in this paper constitutes a useful step towards fully realizing the potential of graph-based references in genomics.

## ACKNOWLEDGMENT

We thank Tony Pan for useful discussions on optimizations presented in this work. We also thank Mehmet Belgin for allocating Stampede2 compute hours for this project. This work was supported by US National Science Foundation grant CCF-1816027 and Extreme Science and Engineering Discovery Environment (XSEDE) resources.

## Footnotes

1 Intel, Xeon and Intel Xeon Phi are trademarks of Intel Corporation or its subsidiaries in the U.S. and/or other countries. Other names and brands may be claimed as the property of others. ©Intel Corporation Software and workloads used in performance tests may have been optimized for performance only on Intel microprocessors. Performance tests, such as SYSmark and MobileMark, are measured using specific computer systems, components, software, operations and functions. Any change to any of those factors may cause the results to vary. You should consult other information and performance tests to assist you in fully evaluating your contemplated purchases, including the performance of that product when combined with other products. For more information go to www.intel.com/benchmarks. Benchmark results were obtained prior to implementation of recent software patches and firmware updates intended to address exploits referred to as “Spectre” and “Meltdown”. Implementation of these updates may make these results inapplicable to your device or system.

## REFERENCES

[1] R. Nielsen, J. S. Paul, A. Albrechtsen, and Y. S. Song, “Genotype and SNP calling from next-generation sequencing data,” Nature Reviews Genetics, vol. 12, no. 6, p. 443, 2011.

[2] A. Dilthey, C. Cox, Z. Iqbal, M. R. Nelson, and G. McVean, “Improved genome inference in the MHC using a population reference graph,” Nature genetics, vol. 47, no. 6, p. 682, 2015.

[3] A. T. Dilthey, P.-A. Gourraud, A. J. Mentzer, N. Cereb, Z. Iqbal, and G. McVean, “High-accuracy HLA type inference from whole-genome sequencing data using population reference graphs,” PLoS computational biology, vol. 12, no. 10, p. e1005151, 2016.

[4] H. P. Eggertsson, H. Jonsson, S. Kristmundsdottir, E. Hjartarson, B. Kehr, G. Masson, F. Zink, K. E. Hjorleifsson, A. Jonasdottir, A. Jonasdottir et al., “Graphtyper enables population-scale genotyping using pangenome graphs,” Nature genetics, vol. 49, no. 11, p. 1654, 2017.

[5] J. A. Sibbesen, L. Maretty, and A. Krogh, “Accurate genotyping across variant classes and lengths using variant graphs,” Nature Publishing Group, Tech. Rep., 2018.

[6] C. P.-G. Consortium, “Computational pan-genomics: status, promises and challenges,” Briefings in Bioinformatics, vol. 19, no. 1, pp. 118–135, 2016.

[7] A. M. Novak, G. Hickey, E. Garrison, S. Blum, A. Connelly, A. Dilthey, J. Eizenga, M. A. S. Elmohamed, S. Guthrie, A. Kahles et al., “Genome graphs,” bioRxiv, 2017.

[8] E. Garrison, J. Sirén, A. M. Novak, G. Hickey, J. M. Eizenga, E. T. Dawson, W. Jones, S. Garg, C. Markello, M. F. Lin et al., “Variation graph toolkit improves read mapping by representing genetic variation in the reference,” Nature biotechnology, 2018.

[9] H. Li and N. Homer, “A survey of sequence alignment algorithms for next-generation sequencing,” Briefings in bioinformatics, vol. 11, no. 5, pp. 473–483, 2010.

[10] J. Sirén, N. Välimäki, and V. Mäkinen, “Indexing graphs for path queries with applications in genome research,” IEEE/ACM Transactions on Computational Biology and Bioinformatics (TCBB), vol. 11, no. 2, pp. 375–388, 2014.

[11] S. Maciuca, C. del Ojo Elias, G. McVean, and Z. Iqbal, “A natural encoding of genetic variation in a burrows-wheeler transform to enable mapping and genome inference,” in International Workshop on Algorithms in Bioinformatics. Springer, 2016, pp. 222–233.

[12] E. Biederstedt, J. C. Oliver, N. F. Hansen, A. Jajoo, N. Dunn, A. Olson, B. Busby, and A. T. Dilthey, “Novograph: Genome graph construction from multiple long-read de novo assemblies,” F1000Research, vol. 7, 2018.

[13] G. Rakocevic, V. Semenyuk, W.-P. Lee, J. Spencer, J. Browning, I. J. Johnson, V. Arsenijevic, J. Nadj, K. Ghose, M. C. Suciu et al., “Fast and accurate genomic analyses using genome graphs,” Nature Publishing Group, Tech. Rep., 2019.

[14] T. F. Smith and M. S. Waterman, “Comparison of biosequences,” Advances in applied mathematics, vol. 2, no. 4, pp. 482–489, 1981.

[15] C. Lee, C. Grasso, and M. F. Sharlow, “Multiple sequence alignment using partial order graphs,” Bioinformatics, vol. 18, no. 3, pp. 452–464, 2002.

[16] G. Navarro, “Improved approximate pattern matching on hypertext,” Theoretical Computer Science, vol. 237, no. 1-2, pp. 455–463, 2000.

[17] R. Vijaya Satya, N. Zavaljevski, and J. Reifman, “A new strategy to reduce allelic bias in RNA-Seq readmapping,” Nucleic acids research, vol. 40, no. 16, pp. e127–e127, 2012.

[18] L. Huang, V. Popic, and S. Batzoglou, “Short read alignment with populations of genomes,” Bioinformatics, vol. 29, no. 13, pp. i361–i370, 2013.

[19] B. Liu, H. Guo, M. Brudno, and Y. Wang, “debga: read alignment with de bruijn graph-based seed and extension,” Bioinformatics, vol. 32, no. 21, pp. 3224–3232, 2016.

[20] D. Kim, J. M. Paggi, and S. Salzberg, “Hisat-genotype: Next generation genomic analysis platform on a personal computer,” bioRxiv, 2018.

[21] J. Pritt, N.-C. Chen, and B. Langmead, “Forge: prioritizing variants for graph genomes,” Genome biology, vol. 19, no. 1, p. 220, 2018.

[22] E. W. Myers and W. Miller, “Approximate matching of regular expressions,” Bulletin of mathematical biology, vol. 51, no. 1, pp. 5–37, 1989.

[23] U. Manber and S. Wu, “Approximate string matching with arbitrary costs for text and hypertext,” in Advances In Structural And Syntactic Pattern Recognition. World Scientific, 1992, pp. 22–33.

[24] A. Amir, M. Lewenstein, and N. Lewenstein, “Hypertext searchinga survey,” in Language, Culture, Computation. Computing-Theory and Technology. Springer, 2014, pp. 364–381.

[25] S. B. Needleman and C. D. Wunsch, “A general method applicable to the search for similarities in the amino acid sequence of two proteins,” Journal of molecular biology, vol. 48, no. 3, pp. 443–453, 1970.

[26] S. Aluru, N. Futamura, and K. Mehrotra, “Parallel biological sequence comparison using prefix computations,” Journal of Parallel and Distributed Computing, vol. 63, no. 3, pp. 264–272, 2003.

[27] Y. Liu, D. L. Maskell, and B. Schmidt, “Cudasw++: optimizing smithwaterman sequence database searches for CUDA-enabled graphics processing units,” BMC research notes, vol. 2, no. 1, p. 73, 2009.

[28] A. Khajeh-Saeed, S. Poole, and J. B. Perot, “Acceleration of the smith– waterman algorithm using single and multiple graphics processors,” Journal of Computational Physics, vol. 229, no. 11, pp. 4247–4258, 2010.

[29] C. Jain and S. Kumar, “Fine-grained GPU parallelization of pairwise local sequence alignment,” in High Performance Computing (HiPC), 2014 21st International Conference on. IEEE, 2014, pp. 1–10.

[30] X. Huang, R. C. Hardison, and W. Miller, “A space-efficient algorithm for local similarities,” Bioinformatics, vol. 6, no. 4, pp. 373–381, 1990.

[31] T. Rognes, “Faster smith-waterman database searches with intersequence SIMD parallelisation,” BMC bioinformatics, vol. 12, no. 1, p. 221, 2011.

[32] S. Misra, T. C. Pan, K. Mahadik, G. Powley, P. N. Vaidya, M. Vasimuddin, and S. Aluru, “Performance extraction and suitability analysis of multi-and many-core architectures for next generation sequencing secondary analysis,” in Proceedings of the 27th International Conference on Parallel Architectures and Compilation Techniques. ACM, 2018, p. 3.

[33] G. P. Consortium, “A global reference for human genetic variation,” Nature, vol. 526, no. 7571, p. 68, 2015.

[34] H. Suzuki and M. Kasahara, “Introducing difference recurrence relations for faster semi-global alignment of long sequences,” BMC bioinformatics, vol. 19, no. 1, p. 45, 2018.

[35] D. S. Hirschberg, “A linear space algorithm for computing maximal common subsequences,” Communications of the ACM, vol. 18, no. 6, pp. 341–343, 1975.

[36] J. A. Grice, R. Hughey, and D. Speck, “Reduced space sequence alignment,” Bioinformatics, vol. 13, no. 1, pp. 45–53, 1997.

[37] M. Holtgrewe, “Mason–a read simulator for second generation sequencing data,” Technical Report FU Berlin, 2010.

[38] Y. Ono, K. Asai, and M. Hamada, “Pbsim: Pacbio reads simulatortoward accurate genome assembly,” Bioinformatics, vol. 29, no. 1, pp. 119–121, 2012.

[39] M. Rautiainen, V. Mäkinen, and T. Marschall, “Bit-parallel sequence-to-graph alignment,” bioRxiv, 2018.

[40] M. Farrar, “Striped smith–waterman speeds database searches six times over other SIMD implementations,” Bioinformatics, vol. 23, no. 2, pp. 156–161, 2006.

[41] R. Vaser, I. Sović, N. Nagarajan, and M. Šikić, “Fast and accurate de novo genome assembly from long uncorrected reads,” Genome research, 2017.

[42] S. Heber, M. Alekseyev, S.-H. Sze, H. Tang, and P. A. Pevzner, “Splicing graphs and EST assembly problem,” Bioinformatics, vol. 18, no. suppl 1, pp. S181–S188, 2002.

[43] W. P. Rowe and M. D. Winn, “Indexed variation graphs for efficient and accurate resistome profiling,” Bioinformatics, vol. 1, p. 8, 2018.

[44] C. Jain, H. Zhang, Y. Gao, and S. Aluru, “On the complexity of sequence to graph alignment,” in International Conference on Research in Computational Molecular Biology. Springer, 2019, pp. 85–100.

